# High-temperature live-cell imaging of cytokinesis, cell motility and cell-cell adhesion in the thermoacidophilic crenarchaeon *Sulfolobus acidocaldarius*

**DOI:** 10.1101/2020.02.16.951772

**Authors:** Arthur Charles-Orszag, Samuel J. Lord, R. Dyche Mullins

**Author notes:** Correspondance to: R.D.M.

## Abstract

Significant technical challenges have limited the study of extremophile cell biology. For example, the absence of methods for performing high-resolution, live-cell imaging at high temperatures (>50°C) has impeded the study of cell motility and cell division in thermophilic archaea such as model organisms from the genus *Sulfolobus*. Here we describe a system for imaging samples at 75°C using high numerical aperture, oil-immersion lenses. With this system we observed and quantified the dynamics of cell division in the model thermoacidophilic crenarchaeon *Sulfolobus acidocaldarius*. In addition, we observed previously undescribed dynamic cell shape changes, cell motility, and cell-cell interactions, shedding significant new light on the high-temperature lifestyle of this organism.

---

Archaea represent one of three domains of life on Earth (Woese *et al.*, 1990), but we know far less about the cell biology of archaeal organisms than we know about bacteria and eukaryotes. For example, we do not understand how most archaea control their shape, organize their intracellular spaces, segregate their DNA, or divide. One reason for this lack of information is that many tools and techniques commonly used to study the cell biology of garden-variety bacteria and eukaryotes do not work properly under the more extreme growth conditions required by some model archaea. This is particularly true for high-resolution, live-cell fluorescence microscopy, which has not been applied to thermophilic prokaryotes mainly because of (i) the difficulty of sufficiently heating samples and high-NA objectives and (ii) a lack of fluorescent proteins that fold correctly and fluoresce at high temperatures.

Despite the lack of live-cell imaging, however, important discoveries regarding chromosome segregation and cell division in crenarchaea have been made in the last decade. These discoveries include the fact that cell scission in many members of the Crenarchaeota is driven, not by homologs of bacterial tubulin-like proteins (i.e. FtsZ), but by relatives of eukaryotic membrane-remodeling proteins with no bacterial homologs.

The genomes of most crenarchaea contain no genes that encode tubulin- or actin-like proteins. In these organisms, daughter cell separation relies on the <u>C</u>ell <u>d</u>i<u>v</u>ision (Cdv) proteins, CdvA, CdvB, and CdvC (Samson *et al.*, 2008; Liu *et al.*, 2017). The CdvB and CdvC proteins are, respectively, homologous to eukaryotic ESCRT-III and Vps4 proteins, which drive membrane remodeling required for: vesicle formation, plasma membrane repair, viral particle budding, and cytokinetic abscission (Liu *et al.*, 2017). In the model crenarchaeon, Sulfolobus acidocaldarius, expression of the CdvABC operon is cell cycle-regulated (Samson *et al.*, 2008). Upon cell division, CdvB, CdvB1, CdvB2 and CdvC assemble a ring-like structure at mid-cell whose diameter decreases as the cytokinetic furrow ingresses (Samson *et al.*, 2008; Risa *et al.*, 2019). It was thus proposed that cell division occurs through constriction of a filamentous CdvB ring previously recruited to the plasma membrane (Dobro *et al.*, 2013). When the ring is first constructed, the CdvB protein inhibits its constriction, but proteasome-mediated degradation of CdvB releases this inhibition, allowing the remaining CdvB1/B2 to constrict (Risa *et al.*, 2019). Evidence from cryoelectrotomograms of dividing cells suggests that the initial phase of ring closure involves only one side of the cell, later extending to the entire cell circumference (Dobro *et al.*, 2013). Although no timelapse imaging has been performed on live, dividing cells, indirect evidence from fluorescence cell sorting suggested that the division process is fast, taking only ~60 seconds to complete (Risa *et al.*, 2019). Direct observation of division, shape change, and motility in live S. *acidocaldarius* will dramatically accelerate research on the fundamental cell biology of crenarchaea.

S. *acidocaldarius* cells are irregular spheroids of 0.8 to 1 μm across, that grow optimally at a temperature of 76°C and at pH 2-3 (Brock *et al.*, 1972). Imaging such small cells under physiologically relevant conditions thus requires (i) high numerical aperture, oil-immersion objective lens, and (ii) a sample chamber capable of maintaining cells at ~76°C. Immersion oil thermally couples the surface of the objective lens to the sample chamber, so to reduce potentially damaging thermal gradients across the optical system and to maintain the cells at a stable temperature, the objective itself must be heated to reduce temperature gradients in the system. Previously, Ogawa and colleagues achieved single molecule imaging of the reverse gyrase from S. tokodaii at 71 °C. In that case, both the imaging chamber and a 100X immersion objective were heated up by circulating hot water through a system of copper tubing connected to a water bath (Ogawa *et al.*, 2015). Here, we describe a microscope system capable of high-resolution imaging live cells at temperatures above 70°C. We use this system to perform time-lapse microscopy of S. *acidocaldarius* cells as they undergo surface motility across glass surfaces; form stable cell-cell contacts; change their shape; and undergo cell division.

## Results

### Microscope system for high-resolution, live-cell imaging at high temperature

We imaged cells by differential interference contrast (DIC) using a Nikon Ti-E inverted microscope equipped with a 1.4 numerical aperture, 100X oil-immersion DIC objective. To maintain cells at 76°C we modified an existing micro-environment control system (DeltaT, Bioptechs), to enable it to achieve higher temperatures than the commercially available version. The sample chamber comprises a 35 mm dish with a heated glass lid and #1.5 glass coverslip coated with an optically transparent but electrically conductive layer of indium tin oxide (ITO). Rather than controlling cell temperature by heating the culture medium or the entire enclosure, our system directly heats the coverglass to which the cells adhere by passing electrical current through the ITO layer via an array of electrodes mounted within the stage adapter (Figure 1A). To maintain cells and culture media at a predetermined temperature, we regulate the electrical current through the ITO layer using a feedback controller and a temperature probe attached to the inner surface of the coverslip. The heated glass lid on top of the sample chamber prevents evaporation of the medium and limits the appearance of advective fluid flows. To reduce thermal loading by the objective lens and to decrease temperature gradients across the metal and glass elements of the optical system, we employed a resistive heater and an additional feedback controller to heat our 100X, 1.4 NA oil-immersion objective. To limit heat transfer to the rest of the microscope we separated the objective lens from the nosepiece by a rubber spacer (Fig. 1A). To determine an appropriate temperature for the objective lens, we measured surface temperatures across the coverslip at various objective temperatures. By heating the objective to 65°C we could maintain an average sample temperature of 75°C with a gradient of 1.1 °C across the field of view (from 74.5°C at the center of the objective to 75.6°C at 5 mm from the center).

**Figure 1.**
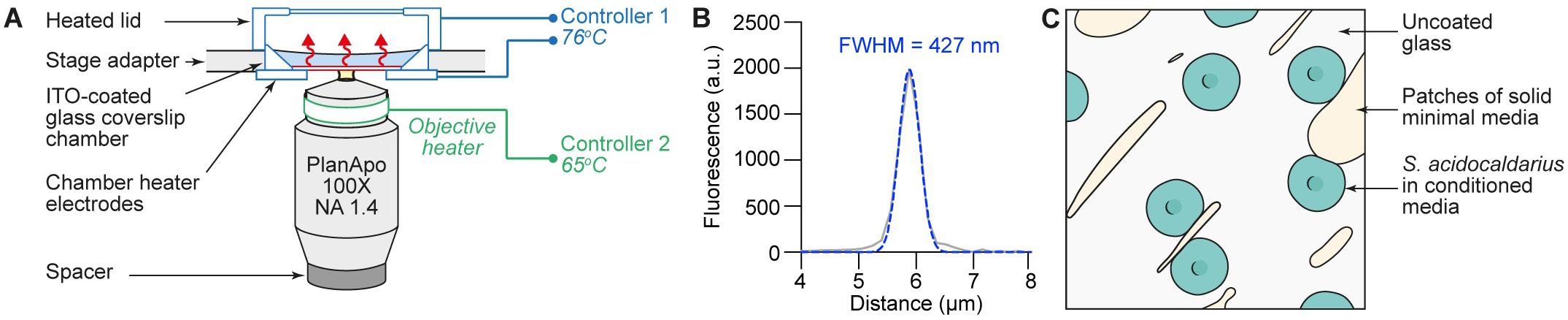
Microscope system for high-resolution live-cell imaging at high temperature. **A.** The Bioptechs DeltaT setup works through an imaging chamber equipped with a glass coverslip whose underside is coated with indium tin oxide (ITO). Passing of a controlled electric current via an array of electrodes through the ITO layer causes the glass surface to generate heat. Evaporation is prevented by a heated glass lid. Another controller maintains a 100X oil immersion objective at 65°C to prevent heat loss from the chamber. **B.** Optical resolution of the system after prolonged imaging at 65/76°C measured with 200 nm fluorescent beads. Grey line: fluorescence intensity; blue dotted line: gaussian fit. **C.** The untreated top side of the glass coverslip is streaked with solid minimal media to enhance cell adhesion. Cells from an exponentially growing culture are diluted in conditioned media.

Happily, no objectives were harmed in the making of this manuscript. To look for evidence of thermal damage to our optical system we measured the point spread function of our 100X 1.4 NA Nikon oil-immersion objective lens after >100 hours of operation at 65°C by imaging 200 nm fluorescent beads at 561 nm illumination (Figure 1B). The two-dimensional point spread function was symmetrical, with a full width at half maximum of 430 nm, which is our minimum resolution. These values were indistinguishable from those before heating the objective.

To characterize cell division under the most physiologically relevant conditions we limited the number of physical and chemical perturbations we applied to the cells. We imaged unsynchronized, wild-type cultures. Experimentally S. *acidocaldarius* cells are synchronized either by treatment with acetic acid (Risa *et al.*, 2019; Lundgren *et al.*, 2004) or use of a microfluidic “baby machine” and subsequent incubation on ice (Duggin *et al.*, 2008; Samson *et al.*, 2008), both of which could potentially perturb the dynamics of subsequent cell divisions. The choice to use unsynchronized populations of cells meant that, to observe the division process, we needed to image cells for long periods of time - significantly longer than the three-hour generation time of this organism at 75°C. To avoid possible constraints on cell motility and perturbations of cell morphology, we also chose not to immobilize cells under a layer of agar (or other gel matrix). When plated on uncoated or poly-L-lysine-coated glass coverslips many cells adhered to the surface but exhibited significant lateral motility and did not stay in the field of view long enough to capture cell division (not shown). After some trial and error we discovered that manual deposition of randomly sized patches (Fig. 1C) of unsupplemented, gelified Brock media (Brock *et al.*, 1972) onto the coverslip provided a set of microenvironments where cells tend to congregate. Other than the patches of gelified media, the coverslip was left untreated. We also decided to image cells sampled from exponentially growing cultures which required a large step dilution of the cells to avoid overcrowding of the field of view. To avoid sending the exponentially growing cells into lag phase, we performed this dilution into conditioned Brock media obtained from a previous exponentially growing culture.

By DIC microscopy most live S. *acidocaldarius* cells growing at 75°C appeared spheroidal, and depending on the focal plane, many of these cells exhibited a central, navel-like depression. Outlines of some cells, however, were noticeably irregular, with sharp vertexes (Fig. 2A). Measure of cells circularity index yielded an average value of 0.92, which better describes a polygon rather than a perfect circle (Fig. 2B). Cell area ranged from 1.2 to 2.7 μm^2^, with an average of 1.7 μm^2^ (Fig 2C). Cells just before division had the same aspect ratio as non-dividing cells (Fig. 2A) with an average circularity index of 0.90 (Fig. 2B). However, cell area was more narrowly distributed around 2 μm^2^, suggesting the existence of a robust cell size control mechanism (Fig 2C).

**Figure 2.**
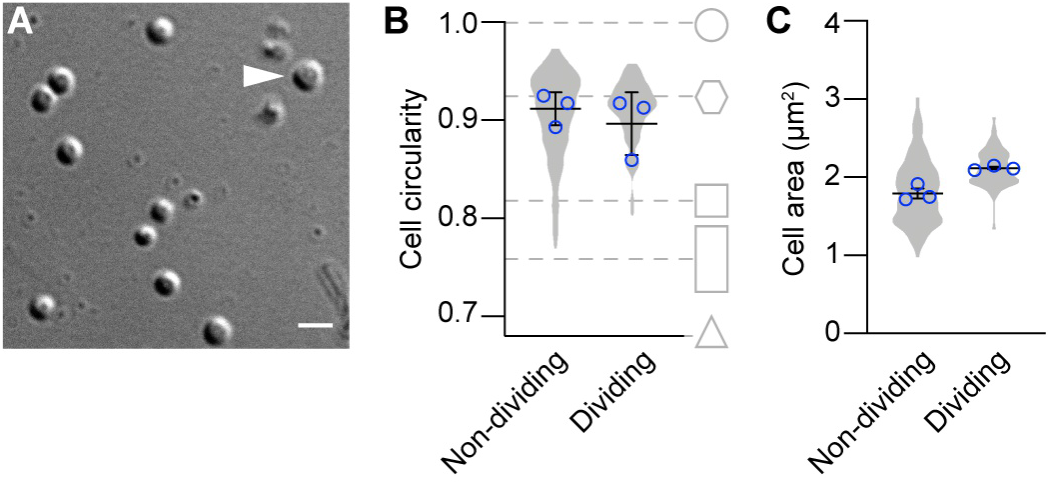
Microscopic characterization of live individual *S. acidocaldarius* cells at high-temperature. **A.** Representative DIC micrograph of cells imaged at 75°C. The arrowhead points to a cell about to divide. Scale bar, 2 μm. **B** and **C.** Circularity index and area of 184 non-dividing and 86 dividing cells. Violin plots correspond to all cells. Blue circles superimposed on the violin plots correspond to the average values in three independent experiments. Bars represent the mean ± SEM.

### Cytokinesis in live S. *acidocaldarius* cells

In more than 10 hours of time-lapse imaging we observed approximately 100 individual examples of cell division in S. *acidocaldarius*, during which one mother cell gave rise to two daughters. The division plane was located at mid-cell. In a significant fraction of cells (20%) ingression of the cleavage furrow began on one side of the cell and progressed unilaterally from one side of the mother cell (Fig. 3A and Movie S1). Ingression otherwise appeared bilaterally symmetrical, although this could reflect an observation angle that masks an underlying asymmetry. Consistent with a central location of the division plane, binary fission most often yielded daughter cells of comparable sizes (Fig. 3B). Rarely (two obvious cases in 86 division events), the division plane was not located at mid-cell and the resulting daughter cells were born with dissimilar sizes (Movie S3, 01:19:00). Time from apparent start to end of division was 116±19 seconds (n=86 cells imaged over 3 experiments, Fig. 3C). Remarkably, the overall shape of dividing cells remained spherical until division was complete. In other words, instead of splitting into two smaller spherical cells, division planes split mother cells into two bean-shaped hemispheres that remained inscribed within the original sphere (Fig. 3D and Movie S2). Many cells exhibited a helical movement relative to one another upon division (Fig. 3E and Movie S3). We did not observe structures reminiscent of a midbody, likely because of the small size of this structure and the important cell movements. When a central navel-like depression could be seen, it was present in each daughter cell immediately after septation. It isn’t clear if this structure disappears and is restored later, or if it is conserved during the whole division process. After division, daughter cells sometimes remained attached to one another by one pole, but generally daughter cells immediately lost adhesion to each other and sprang apart, suggesting a transition from a tensed to a relaxed state. Remarkably, bean-shaped cells relaxed into a spherical shape soon after division was complete (~10-30 seconds).

**Figure 3.**
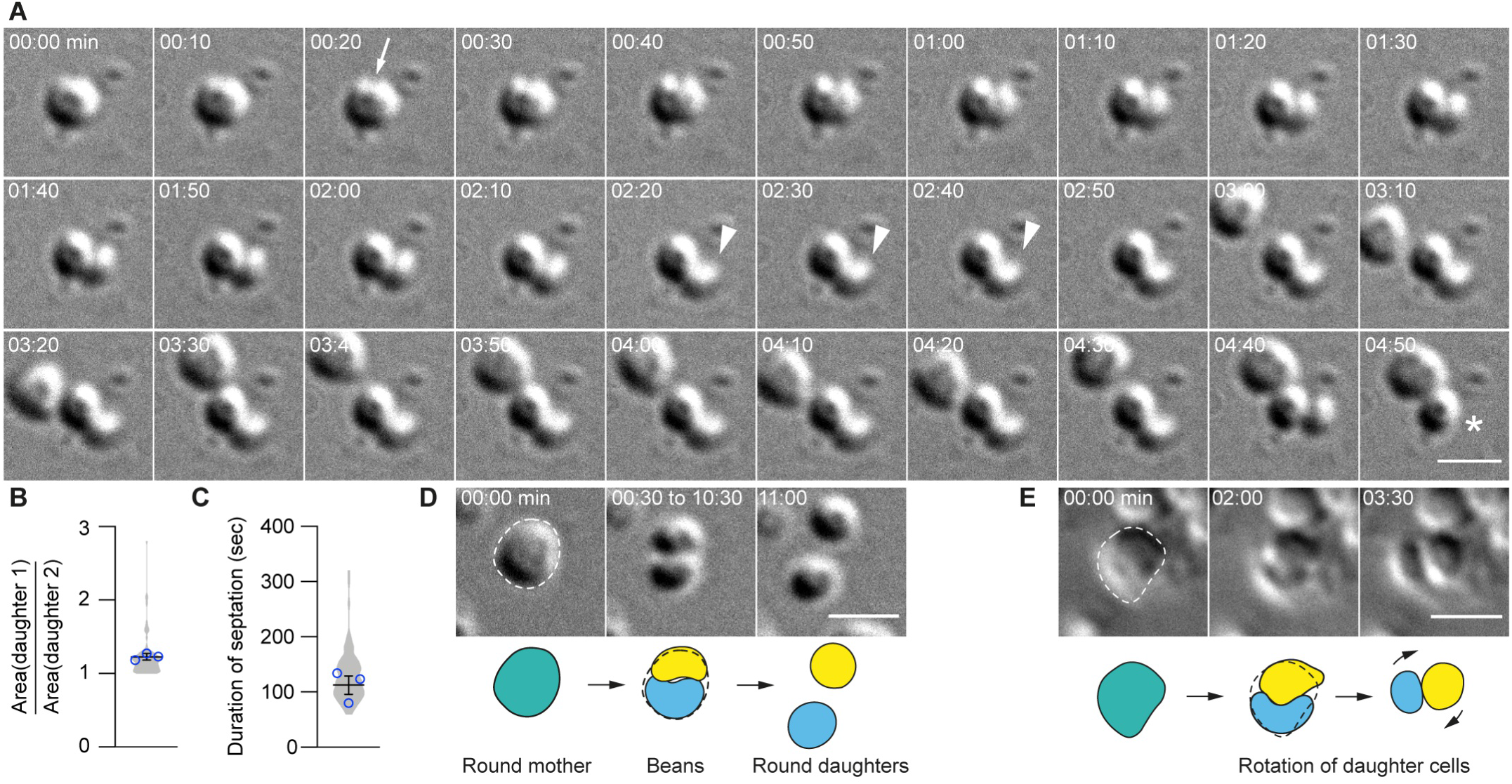
Quantitative live-cell imaging of unsynchronized, unconfined wild-type *S. acidocaldarius* cytokinesis at 75°C. **A.** Snapshots of a representative cell undergoing division (DIC, 150x). The white arrow points to the location where cleavage furrow ingression is initiated. The white arrowheads show the rotation of one daughter cell around its sibling. The asterisks denotes the departure of one of the daughter cells away from its site of birth. See also Movie S1. **B.** Ratio between the areas of the larger and the smaller daughter cell and **C.** apparent duration of division in 86 cytokinesis events. Violin plots correspond to all cells. Superimposed blue circles correspond to the average values in three independent experiments. Bars represent the mean ±SEM. **D.** Representative example of a round mother cell split into two bean-shaped daughters that relax back to a round shape upon separation. See also Movie S2. E. Representative example of a movement of rotation of two daughter cells at birth. See also Movie S3. Interpretative sketches are given at the bottom of D and E. Note that the timepoints are not evenly spaced. Scale bars, 2 μm.

### Surface motility, adhesion and microcolony formation by S. acidocaldarius

In suspension S. *acidocaldarius* cells are very dynamic, likely due to swimming motility (Lewus and Ford, 1999). Interestingly, despite the absence of convective and advective flows in our sample chamber, cells that were adhered to either glass or gelified media exhibited significant lateral motility. Cells movements were often saltatory, reminiscent of type IV pili-mediated twitching motility found in bacteria.

Surface-adhered cells appeared to stop moving prior to cytokinesis. Following this immobile phase, 35±16% of the cells underwent a transient loss of adhesion immediately before dividing, and 45±6% transiently lost surface adhesion upon septation (Fig. 4 and Movie S4). Cells often re-adhered before complete separation of daughter cells. Remarkably, 44±IO% (n=25, N=3) of newly born cells moved rapidly away from their sister cells. In 42±23% (n=26, N=3) of cell divisions, both daughters moved away from the site of division. As a consequence, we did not observe formation of classic microcolonies, as often seen in bacteria.

**Figure 4.**
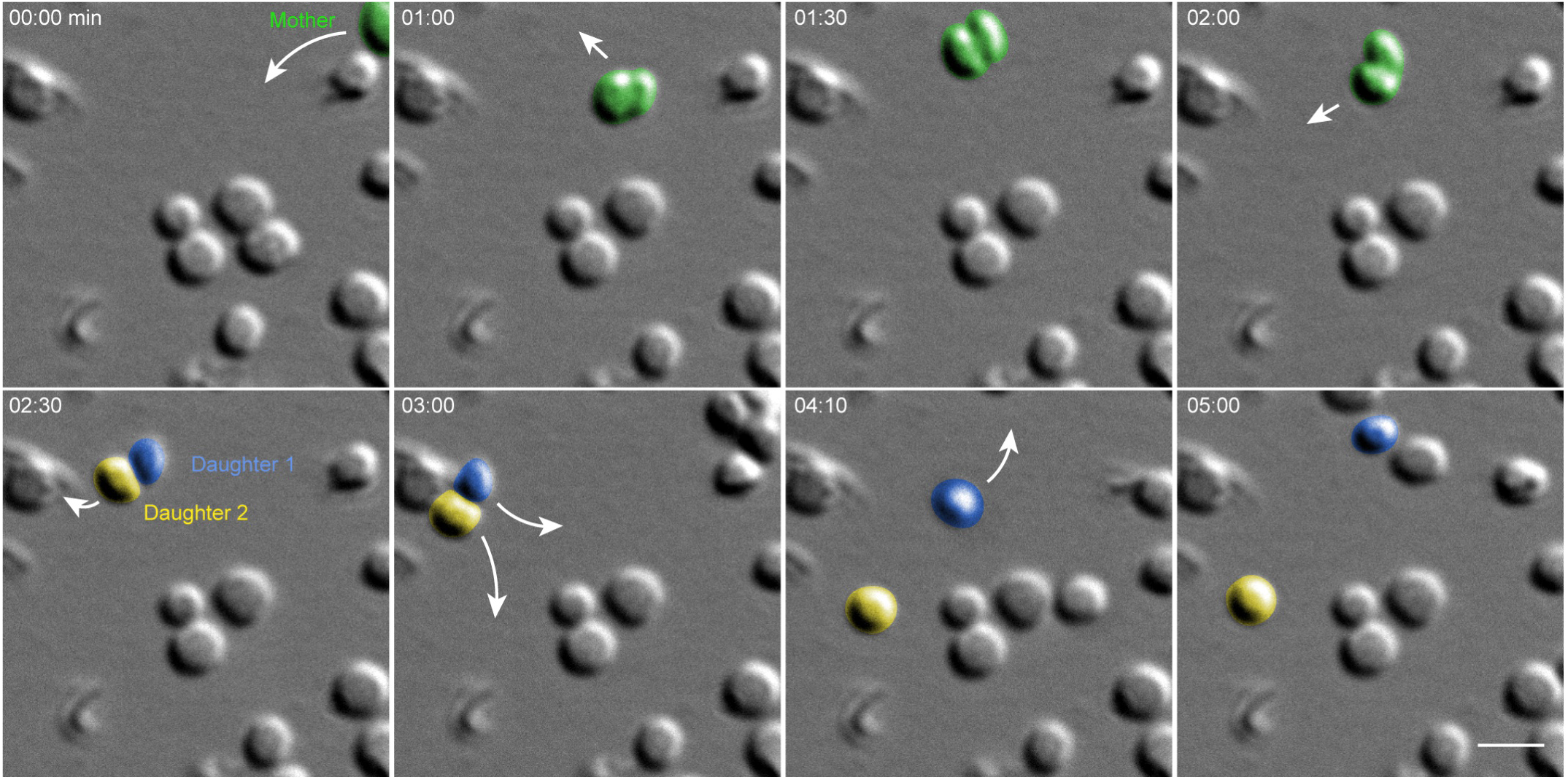
*S. acidocaldarius* cells move around during and after cytokinesis. Snapshots of a representative example of a cell migrating upon and after cytokinesis, producing two free-living daughter cells (DIC, 150x). The white arrows indicate the direction of cell jumps from one site to the next. Note that the timepoints are not evenly spaced. Cells were manually colorized for clarity. See also Movie S4. Scale bar, 2 μm.

### Cell-cell interactions and cell shape transitions in S. *acidocaldarius*

Motile, adherent S. *acidocaldarius* cells exhibited dynamic cell-cell interactions. We observed three distinct behaviors. Firstly, cells were frequently found to interact in pairs. These pairs of cells showed transient “kiss-and-run” adhesion, suggesting specific cell-cell adhesion. These pairwise interactions lasted for prolonged periods of time (114±105 sec, n=92, N=3) before the cells eventually parted (Movie S3). Secondly, cells sometimes switched from a motile to a non-motile state upon encountering a new set of neighbors. Specifically, upon adhesion with two adjacent cells, a motile cell would pause and then attempt to squeeze between two previously attached cells. This behavior produced numerous short chains of polygonal cells resembling rows of cobblestones. In attempting to interpose itself between two adjacent cells, an interloper sometimes moved back and forth between the two neighbors, as if probing its environment and looking specifically for a cell-cell interface. Most surprisingly, this process often required significant dynamic shape changes in the intercalating cell (Fig. 5 and Movie S5). When a cell squeezed between two neighbors in direct contact, the squeezing of the intercalating cell was accompanied by lateral displacement of the neighbors, reflecting either pushing forces exerted by the new cell or active migration of the neighbor cells. Newly intercalated cells remained associated with these “cobblestone chains” for varying lengths of time.

**Figure 5.**
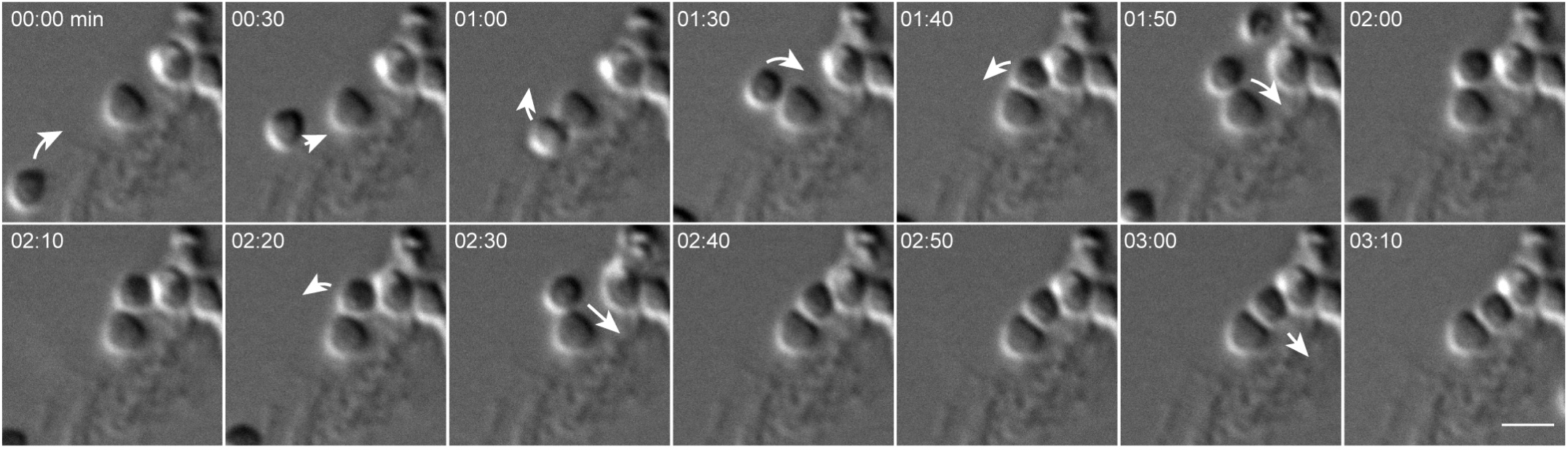
Dynamic cell-cell interactions and squeezing behavior in live *S. acidocaldarius* cells. Snapshots of a representative example of a cell’s behavior upon encoutering new neighbors (DIC, 150x). The white arrows indicate the cell jumps from one site to the next. After migration to a new site, the cell finds a first new neighbor which it seems to interact with. It then finds a gap between the first neighbor and another cell that sits further away and that becomes the second new neighbor. The migrating cell then migrate through the space in between these two new neighbors, thereby changing shape. The process is reversed as the initial cell exits this new space. After a pause, the cell migrates in the space a second time. Again, it then leaves that space to finally come back permanently. Note that the timepoints are not evenly spaced. See also Movie S5. Scale bar, 2 μm.

In addition to shape changes associated with intercalation into pre-existing chains of cells, individual S. *acidocaldarius* cells also exhibited dynamic shape changes during surface associated migration. Cells frequently appeared to flatten at sites of contact with other cells or with the substrate. A representative example of complex cell migration and squeezing behaviors in one cell over multiple hours after birth is shown in Fig. 6 and in Supplementary Movie S6.

**Figure 6.**
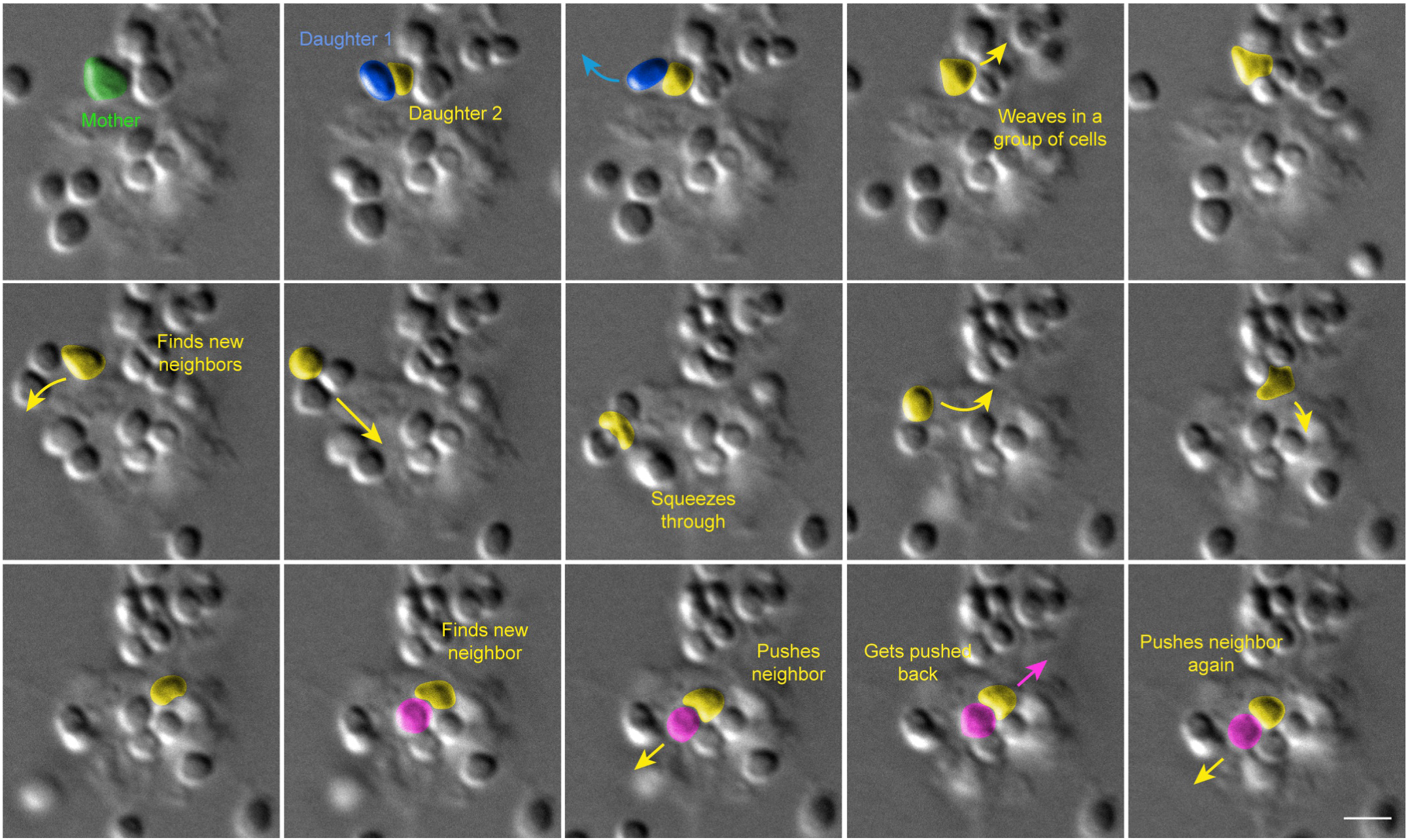
Complex cell shape transitions, migratory and social behaviors in live *S. acidocaldarius* cells over a prolonged period of time. Snapshots of a representative example of a cell’s complex life over an hour after its birth (DIC, 150x). The arrows indicate the cell jumps from one site to the next. Text indicates important steps. Note that the snapshots were voluntarily discontinued in time to highlight important frames. Cells were manually colorized for clarity. Note that the timepoints are not evenly spaced. See also Movie S6. Scale bar, 2 μm.

## Discussion

At some point every microscopist is tempted to deploy the famous Yogi Berra quote: “You can observe a lot by just watching.” In this case, we observed a lot of new phenomena simply by performing high-resolution time-lapse imaging of cells that had never been imaged at physiologically relevant temperatures. For this work we created and validated a system for long-duration (>4 hours), high-resolution, time-lapse imaging of live S. *acidocaldarius* cells at high temperature (up to 75°C). In addition to the fact that we observed surface motility and cell division, our results are quantitatively consistent with previous studies, including light and electron microscopy of fixed cells. Specifically, the following measured values are consistent with previous literature: (i) cell size and aspect ratio (Brock *et al.*, 1972); (ii) generation time (Duggin *et al.*, 2008); and (iii) time required for cytokinesis (Risa *et al.*, 2019). Specifically, we find that cytokinesis in these cells requires 116 seconds, which turns out to be 15 to 30 times faster than cytokinesis in bacteria such as *E. coli*. The best characterized examples of bacterial cytokinesis are driven by the activity of peptidoglycan synthetases, organized into rings by tubulin-like polymers (e.g. FtsZ), which require 30 to 60 minutes to form a septum separating daughter cells in bacteria such as *E. coli* (Coltharp *et al.*, 2016), S. *aureus* (Monteiro *et al.*, 2015), and *C. crescentus* (Li *et al.*, 2009). A similar mechanism appears to drive cell division in the halophilic euryarchaeon *H. volcanii* (Walsh *et al.*, 2019). Our data agree well with the prediction of Risa, Hurtig and colleagues that the duration of cytokinesis in S. *acidocaldarius* is ~6O seconds (Risa *et al.*, 2019). Further experimentation will be required to determine whether cell division is as fast in other crenarchaea that also employ the CdvAB system.

Our work also verifies in live cells a proposal of Risa, Hurtig *et al.* (2019) regarding the geometry of dividing cells. Based on simulations of ESCRT-III-mediated division of nearly spherical cells Risa, Hurtig *et al.* (2019) suggested that daughter cells adopt “bean” shapes inscribed within the sphere defined by the mother cell. Our time-lapse imaging revealed that dividing cells adopt this overall geometry just before daughter cells snap apart. We did not, however, detect a gap between the two opposite sides of the cleavage furrow, and we found that sister cell remained in close contact until septation was complete. At the end of septation daughter cells rapidly detach from each other, often with a helical or twisting motion (Fig. 3E and Movie S3). Cell shape then quickly (within 30 seconds) relaxes from hemispherical to approximately spherical. These observations suggest that constriction of the ESCRT-III ring generates torsional stress at the division site that is released upon separation of the daughter cells in a spring-like fashion. We speculate that this behavior might reflect inhomogeneities in the distribution of S-layer proteins at the end of cytokinesis. Specifically, the rapid ingression of the cleavage furrow might result in the absence of S-layer proteins on this region of the membrane. Subsequent reorganization of S-layer proteins and/or catastrophic detachment of S-layer connections around the circumference of the original mother cell might contribute to the abrupt movement of daughter cells. Additional work will be required to understand the dynamics of S-layer assembly and reorganization during and after cell division and whether they play a role in the cell movements we observe.

Our data also suggest that S. *acidocaldarius* is capable of twitching motility similar to that driven by retractile pili in bacteria (e.g. *N. meningitidis*). The genome of S. *acidocaldarius* encodes three different types of filaments related to type IV bacterial pili: archaella, which power swimming motility; UV-inducible pili, which promote exchange of DNA through autoaggregation upon UV-induced DNA damage; and archaeal adhesive pili (Aap), which mediate surface adhesion (Henche *et al.*, 2012a; Henche *et al.*, 2012b). In bacteria, type IV pili can drive twitching motility, characterized by saltatory, lateral movements across a surface. These movements are generated by the disassembly of surface-attached pili under the action of the retraction ATPase PilT (Merz *et al.*, 2000). However, although homologs of the assembly ATPase PilF are found in crenarchaeal genomes, there are no homologs of PilT (Berry and Pelicic, 2015), raising the question of whether S. *acidocaldarius* adhesive pili can generate force via retraction. Intriguingly, Ellison *et al.* (2017) recently showed that type IV pili of the bacterium *C. crescentus*, which lacks homologs of PilT can, nevertheless, undergo retraction (Ellison *et al.*, 2017). These authors found that, in C. *crescentus*, a unique ATPase, CpaF, mediates both elongation and retraction of the pili (Ellison *et al.*, 2019), and we speculate that a similar PilT-independent mechanism might drive type IV pilus retraction in S. *acidocaldarius*, thus enabling twitching motility. Given the lack of direct evidence that the cells express adhesive pili under the experimental conditions tested here, further work will be required to directly assess whether mutants defective for pilus biogenesis are capable of surface motility.

In addition to the surface motility, we were surprised by the dynamic nature of substrate adhesion and cell-cell interactions in S. *acidocaldarius*. In particular, we were impressed by the fact that (i) the large majority (87.5%) of newly divided daughter cells lose substrate attachment at the moment of scission; and (ii) chains of attached cells are formed when individual motile cells encounter each other or preformed chains and settle down to make stable attachments. These observations have several implications for microcolony assembly and biofilm formation. In bacteria, cells frequently form microcolonies and biofilms by remaining attached to their sisters following division. As cells divide and the colony grows bigger, the secretion of extracellular polymeric substances (EPS) and extracellular DNA from dead cells collaborate to form a biofilm matrix. The mature biofilm provides the cell community with protection against external challenges such as antimicrobials, and promotes cell survival through enhanced nutrient capture (Flemming *et al.*, 2016). A notable exception to this is the bacterium *C. crescentus*. Upon adhesion to the substrate, cells differentiate into stalked cells that are incapable of swimming. Cell division occurs in an asymmetrical way and results in the release of a motile swarmer cell from the stalked cell. The same stalked cell can undergo multiple rounds of division, each yielding one swarmer cell capable of colonizing distant sites but incapable of division. This lifestyle is thought to promote survival in nutrient-deprived environments such as the natural habitat of C. *crescentus* (Woldemeskel and Goley, 2017). However, this species is still capable of biofilm formation, suggesting either canonical microcolony formation under nutrient-rich conditions or differentiation of swarmer cells in the vicinity of new stalker cells long after birth. The same survival strategy could be used in S. *acidocaldarius* which inhabits nutrient-poor hot springs. This would be consistent with reports in the literature that, although S. *acidocaldarius* forms biofilms, microcolony formation occurs late in the organism’s life cycle. Scanning electron microscopy revealed that microcolonies are detectable only after around 36h *in vitro* (Henche *et al.*, 2012b), which would be consistent with microcolony formation driven by motility and congregation of non-sister cells rather than by persistent attachment of sisters. Further experiments comparing the behavior of *S. acidocaldarius* cells under various nutrient availability conditions will be required.

We noticed that the intercalation of individual S. *acidocaldarius* into previously formed cell chains was accompanied by dramatic cell shape changes. To our knowledge, this phenomenon has not been previously reported in either bacteria or archaea. These observations suggest the existence of cell-cell adhesion molecules and machinery for generating extra- and/or intracellular forces capable of deforming the cell. Cells that squeezed between two existing neighbors appeared to actively pull themselves into the cleft, pushing the neighbors apart, in a diapedesis-like fashion. Since S. *acidocaldarius* lacks homologs of actin- and tubulin-like proteins, and since ESCRT-III homologs mediate large cell surface rearrangements upon cell division, it is tempting to hypothesize that the ESCRT-III machinery might mediate such cell shape changes. The expression of ESCRT-III genes, however, is tightly linked to the cell cycle in S. *acidocaldarius* (Samson *et al.*, 2008), implying either that cells are competent for shape transitions only during the G2/M phase or that other systems control cells shape throughout the cell cycle. Both monoderm and diderm bacteria possess a rigid peptidoglycan-based cell wall which resists large scale plastic deformations. Archaeal cell walls, however, are very different. In S. *acidocaldarius*, the plasma membrane is made of a monolayer of tetraether lipids, topped with a paracrystalline array of SlaA and SlaB proteins, which form the S-layer (Gambelli *et al.*, 2019; Rodrigues-Oliveira *et al.*, 2017). The S-layer was recently shown to protect cells of a related archaeal species, S. *islandicus*, against osmotic stress (Zhang *et al.*, 2019). In S. *solfatarícus* the S-layer protects cells against viral infection and plays an important role in cell division (Zink *et al.*, 2019). The deformability of the S-layer has not been measured *in vivo*, but links have been proposed between S-layer architecture and cell shape. For example, Taylor *et al.* (1982) suggested that different arrangements of SlaAB arrays might account for various structural features on the cell surface, such as vertices and lobes (Taylor *et al.*, 1982).

Evidence for intimate cell-cell contacts has been observed in scanning electron micrographs of early biofilm formation in S. *acidocaldarius* (Koerdt *et al.*, 2010). However, even if UV-inducible pili promote autoaggregation, we did not observe such aggregates under the experimental conditions tested, consistent with the fact that we used relatively long wavelength (520-530 nm) light to image the cells. This result suggests that adhesion molecules other than UV-inducible pilins can promote cell-cell contact in S. *acidocaldarius*. Interestingly, multiple genes in the S. *acidocaldarius* genome share homology with cell-cell adhesion molecules from other species, including bacterial antigen 43 of *E. coli (saci_2215)*, the serine-rich α-agglutinin of *Saccharomyces cerevisiae (saci_2214* and *saci_1140)*, and filamin repeat-containing eukaryotic adhesion proteins *(saci_2356)*. These genes are excellent candidates to mediate the type of cell-cell contacts we observe at 75°C in S. *acidocaldarius*.

Our high-temperature microscope can be adapted and or improved in several ways. Firstly, employing microchambers and synchronized cell cultures would increase the rate of capturing cytokinesis events and enable us to follow the same cell over multiple division events. This would help determine the mechanism underlying cell size control. Secondly, the development of high-temperature fluorescent proteins that fold correctly in archaeal cells (Frenzel *et al.*, 2018; Scott *et al.*, 2018; Wannier *et al.*, 2018) combined with the genetic tractability of Sulfolobales (Wagner *et al.*, 2012) would dramatically improve our ability to address molecular mechanisms underlying cellular behavior.

## Materials and methods

### Cell culture and preparation of conditioned media

The wild type Sulfolobus acidocaldarius strain DSM 639 (generous gift from Sonja-Verena Albers, University of Freiburg) was grown aerobically at 75°C in Brock’s minimal media (Brock *et al.*, 1972) supplemented with 0.1% tryptone (Sigma-Aldrich).

Conditioned Brock’s media was prepared by centrifugation of 50 mL of an exponentially growing culture (OD_600nm_ ~0.2-0.3) at 4,000 *×g* and room temperature for 15 min followed by syringe-filtration through a 0.22 μm filter to remove cells and cell debris. Conditioned media was distributed in 15 mL conical tubes and stored at 4°C for up to a month.

### High-resolution imaging at high temperature

A DeltaT cell micro-environment control system (Bioptechs) was modified by the manufacturer to allow it to reach temperatures up to 80°C. It was used on a Nikon Ti-E inverted microscope equipped with a motorized stage.

In order to enhance cell attachment, 2X Brock’s minimal media at 75°C was mixed with an equal volume of freshly boiled 1.7% gelrite (Gelzan, Sigma). A sterile pipet tip was used to manually streak the bottom coverslip of a DeltaT imaging chamber with that solution. The media was allowed to solidify for 5 min at room temperature and 2 mL of conditioned Brock’s media were added to the chamber. Modified chambers were prepared the day of the experiment and kept in an incubator at 75°C until imaging. The microscope objective was prewarmed to 65°C prior to imaging and kept at this temperature throughout the experiment. Upon imaging, 500 μL of cells were sampled from an exponentially growing culture (OD_600nm_ ~0.2-0.3), added to an imaging chamber and immediately set on the microscope stage to avoid prolonged drop in temperature. The chamber controller was set to 78°C to maintain cells and media between 74.5 and 75.6°C throughout the experiment. The same controller was used heat a glass lid that which temperature was manually increased until no condensation was observed.

Differential interference contrast (DIC) microscopy was performed with a green light-emitting diode illuminator (B180-RGB; ScopeLED) through a Nikon Plan Apo VC 100X 1.4 NA oil objective with an additional 1.5x tube lens, and a Point Grey CMOS camera (CM3-U3-50S5M-CS; FLIR). Cells were imaged with a 100-ms exposure every 10 second. All microscopy hardware was controlled with Micro-Manager software (Edelstein *et al.*, 2010).

### Data analysis

Image analysis and quantifications were manually made in Fiji (ImageJ (Schindelin *et al.*, 2012)). Data was analyzed and graphs were prepared in Prism (GraphPad). Figures and illustrations were prepared in Adobe Illustrator.

## Acknowledgments

We would like to thank Sonja-Verena Albers for sharing the cells, Buzz Baum, Deylan Mutavchief and Andre Arashiro Pulschen for fruitful discussions, and Joshua Edwards for initial help with the imaging setup. This work was supported by the National Institute of General Medical Sciences of the National Institutes of Health (R01-GM061010 to R.D. Mullins), by the Howard Hughes Medical Institute Investigator program (R.D. Mullins, A. Charles-Orszag). The authors declare no competing financial interests.

## Author contributions

A. Charles-Orszag: conceptualization, methodology, investigation, formal analysis, validation, visualization, writing (original draft preparation), and writing (review and editing). S.J. Lord: methodology, investigation, formal analysis, validation, visualization, figure preparation, writing (original draft preparation), and writing (review and editing). R.D. Mullins: funding acquisition, supervision, methodology, investigation, formal analysis, and writing (original draft preparation, review, and editing).

